# Episodic memory supports the acquisition of structured task representations

**DOI:** 10.1101/2024.05.06.592749

**Authors:** Qihong Lu, Ali Hummos, Kenneth A. Norman

## Abstract

Generalization to new tasks requires learning of task representations that accurately reflect the similarity structure of the task space. Here, we argue that episodic memory (EM) plays an essential role in this process by stabilizing task representations, thereby supporting the accumulation of structured knowledge. We demonstrate this using a neural network model that infers task representations that minimize the current task’s objective function; crucially, the model can retrieve previously encoded task representations from EM and use these to initialize the task inference process. With EM, the model succeeds in learning the underlying task structure; without EM, task representations drift and the network fails to learn the structure. We further show that EM errors can support structure learning by promoting the activation of similar task representations in tasks with similar sensory inputs. Overall, this model provides a novel account of how EM supports the acquisition of structured task representations.

## Reusing task representations via EM

Prior neural network modeling work has shown that having an explicit representation of the current task (in a separate model layer, projecting to the rest of the model) can facilitate task performance by reducing interference from other tasks (Flesch, Nagy, Saxe, & Summerfield, 2023; Masse, Grant, & Freedman, 2018; Lu et al., 2023; Russin, Zolfaghar, Park, Boorman, & O’Reilly, 2022). Crucially, this representation should also reflect the structure *across* related tasks, so knowledge about other tasks can be leveraged for generalization to new tasks (Yang, Joglekar, Song, Newsome, & Wang, 2019; Flesch, Saxe, & Summerfield, 2023; Ito et al., 2022; Tafazoli et al., 2024). Consider the problem of driving. Vehicles travel on the right in the US and on the left in the UK. When traveling from the US to the UK for the first time, one needs to construct a UK representation to avoid interference, but this representation also needs to be structurally related to the US representation to benefit from prior knowledge about driving. How should an intelligent agent build task representations online that satisfies both criteria?

Previous studies have suggested that episodic memory (EM) contributes to this process by retrieving previously formed task representations. This claim is supported by a large body of experimental work (for a review, see Egner, 2023). It also resonates with theoretical work on *amortized inference*, which posits that inferring an appropriate task representation online can be computationally costly, and that performance can be improved by re-using the products of these costly inference processes (Dasgupta, Schulz, Goodman, & Gershman, 2018; Dasgupta & Gershman, 2021); in line with this view, it has been shown that EM in particular can support efficient re-use when the same task repeats (Dasgupta et al., 2018; Lu et al., 2023).

Here, we go beyond prior work by demonstrating that EM helps the model learn task representations that veridically reflect the structure *across* tasks. This contribution of EM to structure learning arises for two reasons: First, inferring a task representation with information about the ongoing task only is underdetermined – representations that support good performance for the ongoing task might not be unique, causing the representation for a task to drift over time. This lack of stability can hinder the accumulation of relational structure across tasks – it is difficult to build an intricately structured building on shifting sands. EM can counteract this instability by biasing the network to consistently re-use previously discovered task representations. We also demonstrate that the stabilizing influence of EM is sometimes not enough to support structure learning, but in this case EM *errors* can help to rescue structure learning by promoting activation of similar task representations in tasks with similar sensory inputs.

## Model architecture

We developed a neural network augmented by a task representation layer and an EM buffer (Fig. 1a). The model infers a task representation at each task switch point (task switches are explicitly signaled to the model – a simplification that allows us to focus on the influence of EM on task representation). Several recent models of task inference have assumed orthogonal or non-overlapping task representations to i) avoid interference across tasks and ii) improve computational feasibility (Franklin, Norman, Ranganath, Zacks, & Gershman, 2020; Lu et al., 2023); as a result, these models can not directly represent similarity structure across tasks. To get around this limitation, we leveraged the idea of using gradient-based search in activation space to infer task representations online (Giallanza, Campbell, Cohen, & Rogers, 2023; Hummos, 2022): After a task switch, the model finds a task representation by adjusting the task layer activation to optimize the ongoing task objective. Concretely, the model collects a small sample of observations from the ongoing task. Then, to minimize the loss of this sample, the model performs gradient descent in the space of task representations until convergence (Fig. 1b), while holding network weights fixed. The discovered task representation is held constant within the ongoing task (i.e., until the next task switch occurs; Flesch, Nagy, et al., 2023; Russin et al., 2022). As the task representations are not constrained to be orthogonal, this model has greater flexibility for learning rich relations across tasks. We focus here on blocked (rather than interleaved) curricula, which are challenging for standard neural networks due to catastrophic interference (McClelland, McNaughton, & O’Reilly, 1995).

**Figure 1:**
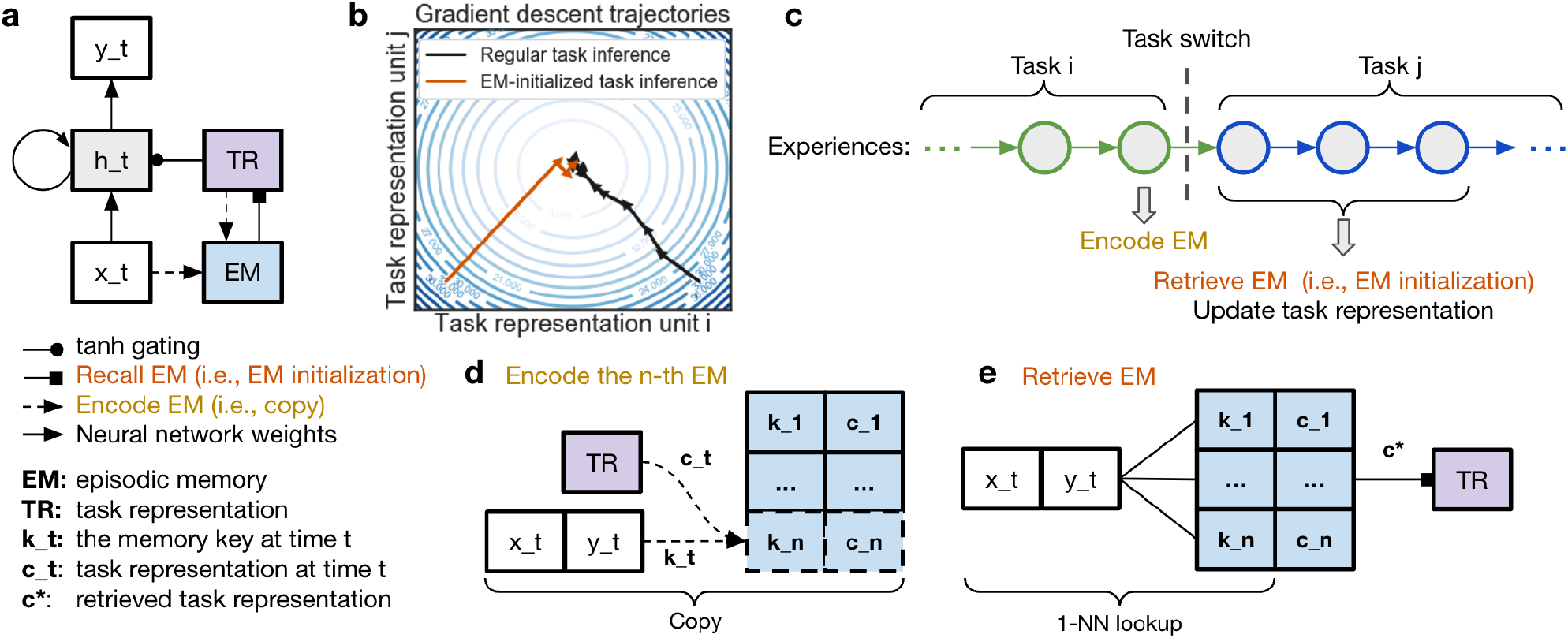
**(a)** A neural network model with a task representation (TR) layer and an episodic memory (EM) module that encodes and retrieves task representations. **(b)** The model infers a task representation by performing gradient descent until convergence in the space of task representations (activation patterns) to minimize the loss objective of the ongoing task; during this inference process, the network weights are not adjusted. The inferred task representation is a locally optimal solution to the current task objective. EM can speed up this process by retrieving a previously encoded task representation to *initialize* the gradient search. **(c)** At the end of every task, the model encodes an EM. After a task switch, the model collects a small sample of observations from the ongoing task. This small sample is first used as a cue to retrieve a previously encoded EM, which initializes the task inference process. Then, the gradient search process is executed to optimize the task objective on this small sample. **(d)** To encode a new memory (the n-th memory in this demo), the model first uses the current observation *{x*_*t*_, *y*_*t*_*}* to form a memory key, *k*_*t*_. Then, the model simply stores a copy of the current task representation, *c*_*t*_, indexed by *k*_*t*_. **(e)** To recall a memory at time t, the model uses the current observation *{x*_*t*_, *y*_*t*_*}* to form a memory key *k*_*t*_, and performs a one-nearest-neighbor search for the task representation *c*^*\**^ with the most similar memory key.

Our model is augmented with a non-differentiable EM module (Fig. 1a; Pritzel et al., 2017; Ritter et al., 2018). EM encoding occurs right before every task switch (Fig. 1c). Each memory encodes i) the task representation, *c*_*t*_, and ii) a memory key *k*_*t*_ for memory indexing (Fig. 1d). The memory key is simply a copy of the sensory experience *x*_*t*_ and *y*_*t*_. Our assumption that EM encoding selectively occurs at the ends of events is consistent with empirical data (Baldassano et al., 2017; Ben-Yakov & Henson, 2018; Barnett et al., 2023); prior modeling work has shown that this is computationally optimal for retrieval success (Lu, Hasson, & Norman, 2022).

When learning temporal sequences (see Simulation 1), the model replays the just-experienced sequence at the end of the task to obtain a more informative task representation, which is then stored in EM. Concretely, replay involves performing task inference by optimizing the objective over the entire just-experienced sequence rather than a limited sample of initial observations. Without replay, the task representation can overfit the observations available at the beginning of that task.

Episodic retrieval is similarity-based and occurs after a task switch (Fig. 1c). First, the model forms a memory key, *k*_*t*_, using the current observations. Then, it performs a one-nearest-neighbor search over all memories (Fig. 1e) to find the previously formed task representation with the most similar memory key. The retrieved task representation initializes the task inference process. Finally, the model refines this retrieved task representation by performing task inference (i.e., gradient-based optimization of task representation to optimize the ongoing objective). This refinement is important, as a retrieved representation stored in the past might not be aligned with the current network weights (which are constantly adjusted for learning purposes).

## Simulation 1. Learning compositional structure

We developed a task-dependent sequence-learning environment (Fig. 2a) based on the paradigm from Beukers et al. (2023). The task varies along three orthogonal dimensions. All dimensions are binary – the first dimension is either café (0) or bar (1); the second dimension is either chat (0) or work (1); the third dimension is either jazz (0) or blues (1). Every task dimension influences the transition structure across states independently. For example (see Fig. 2b), if the current task is bar + work + jazz, then state [1, 4, 5] leads to [7, 10, 12].

**Figure 2:**
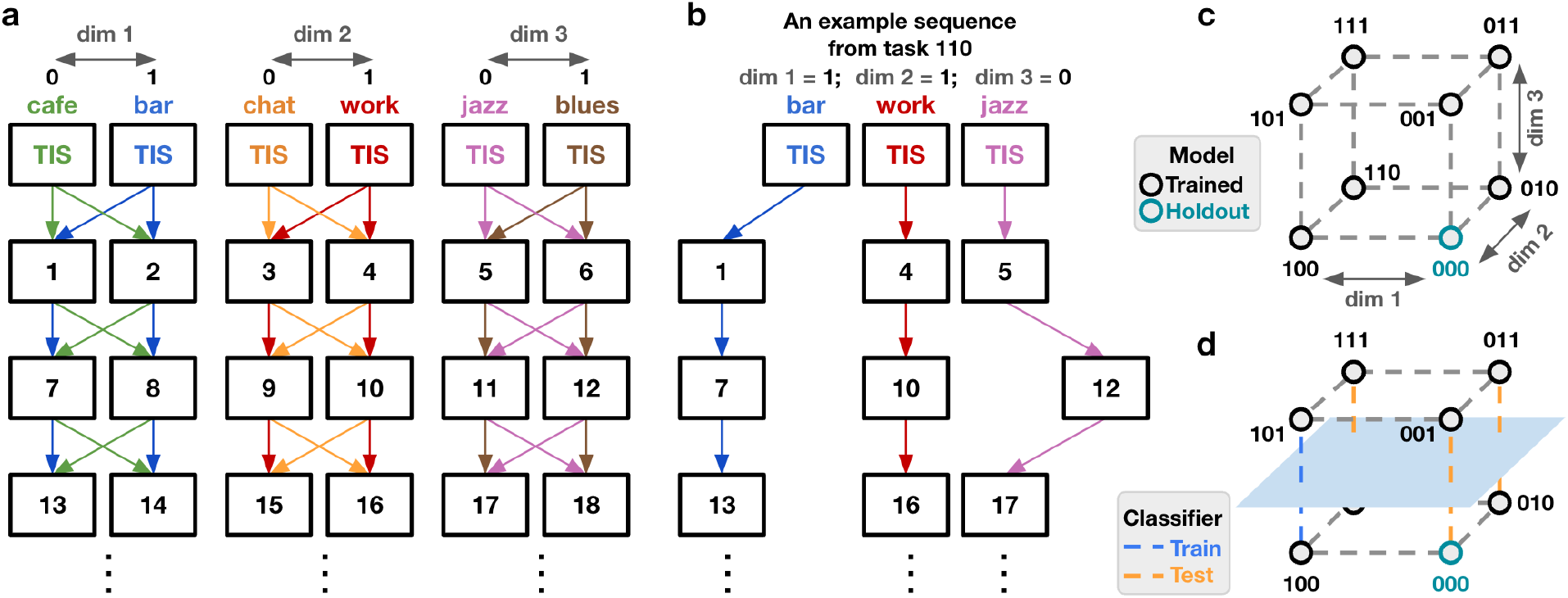
**(a)** The graph used to generate task-dependent sequences in Simulation 1. State transitions are controlled by the task identity (e.g., bar + work + jazz). For example, in a bar, the blue transitions will be used. At the very beginning of the sequence, a three-dimensional task-indicative signal (TIS) is presented. **(b)** In the bar + work + jazz task (i.e., task 110), the state [1, 4, 5] goes to [7, 10, 12], and then [13, 16, 17]. **(c)** The ideal neural representational geometry for the eight tasks should form a cube, as the three binary task dimensions are orthogonal. **(d)** To test if the representation for the third dimension is orthogonal to the other two, we can compute the cross-condition generalization performance (CCGP; Bernardi et al., 2020) along dimension 3: We can start with training a classifier to differentiate the representations for task 101 versus 100. Then, we test the resulting decision plane (marked in light blue) to differentiate the third dimension while varying the other two task dimensions (i.e., 001 versus 000, 111 versus 110, and 011 versus 010). If the test accuracy is 1 for all combinations of train-test splits and across all three dimensions, then CCGP is maximal (i.e., 1).

Note that, if the first task dimension changes from bar to cafe (i.e., in a cafe + work + jazz task), then [1, 4, 5] leads to [**8**, 10, 12] – only the first element of the state is impacted.

Given the current state, the model’s objective is to predict the upcoming state. At the beginning of every sequence, a three-dimensional task-indicative signal (TIS) was presented (e.g. the TIS for bar + work + jazz is 110, see Fig. 2b). All models were trained on a blocked curriculum for six epochs. In every epoch, the model cycles in a random order through 7 of the 8 tasks (all tasks except for the holdout task 000; Fig. 2c), receiving a block of 200 sequences per task.

At the end of every sequence, the model first replays the just-experienced sequence to update its task representation. Then, it encodes a memory consisting of the TIS as the memory key and the current task representation. Whenever there is an existing memory indexed by the same memory key, the encoded task representation for that old memory is updated using the new one. Then, the weights are updated to minimize the loss of this sequence. At the beginning of every new sequence, the model first uses the just-observed TIS to retrieve a previously formed task representation to initialize task inference. Then, the model performs task inference using the initial observations of the sequence.

Ideally, the neural representations for the eight tasks should form a cube (Fig. 2c). In other words, the coding direction representing the cafe-bar difference should be “abstract” (as defined by Behrens et al., 2018; Bernardi et al., 2020; Vaidya & Badre, 2022) – it should be invariant to whether the current task is chat or work and jazz or blues. We used cross-condition generalization performance (CCGP; Bernardi et al., 2020) to measure abstraction along the three task dimensions. For example, consider dimension 3: We can start with training a classifier to differentiate the neural representations for task 000 versus 001. This classifier should generalize regardless of what the first two task dimensions are. Namely, the resulting decision plane (the light blue plane in Fig. 2d) should also reliably classify 100 versus 101, 010 versus 011, and 110 versus 111. CCGP is maximal (i.e., 1) if the classifier generalization performance is 1 for all three task dimensions.

To understand the effect of EM, we trained three kinds of models: Our baseline was a recurrent neural network (RNN) with Gated Recurrent Units (*GRU*; Cho et al., 2014) but no task layer and no EM; we compared this to a *no-EM* model (a GRU with a task layer but no EM) and a *with-EM* model (a GRU with a task layer an EM). Then we measured performance during training (Fig. 3b) and test (Fig. 3c) as well as CCGP for the learned task representations during test (Fig. 3a). During training, the (baseline) GRU was slow at learning (Fig. 3b) as blocked training led to catastrophic interference across tasks (McClelland et al., 1995). The no-EM model showed a distinctive pattern of performance where its performance was poor at the start of task blocks but it improved sharply within blocks. The improvement within blocks can be explained in terms of the model inferring task representations that help it minimize across-task inference; the drop at block boundaries can be explained in terms of the model failing to re-find old task representations during the inference process, which impedes the accumulation of knowledge. With EM, the performance reached the ceiling almost immediately after a task switch (Fig. 3b), as the model was able to re-load previously used task representations that give it access to knowledge acquired in previous blocks.

**Figure 3:**
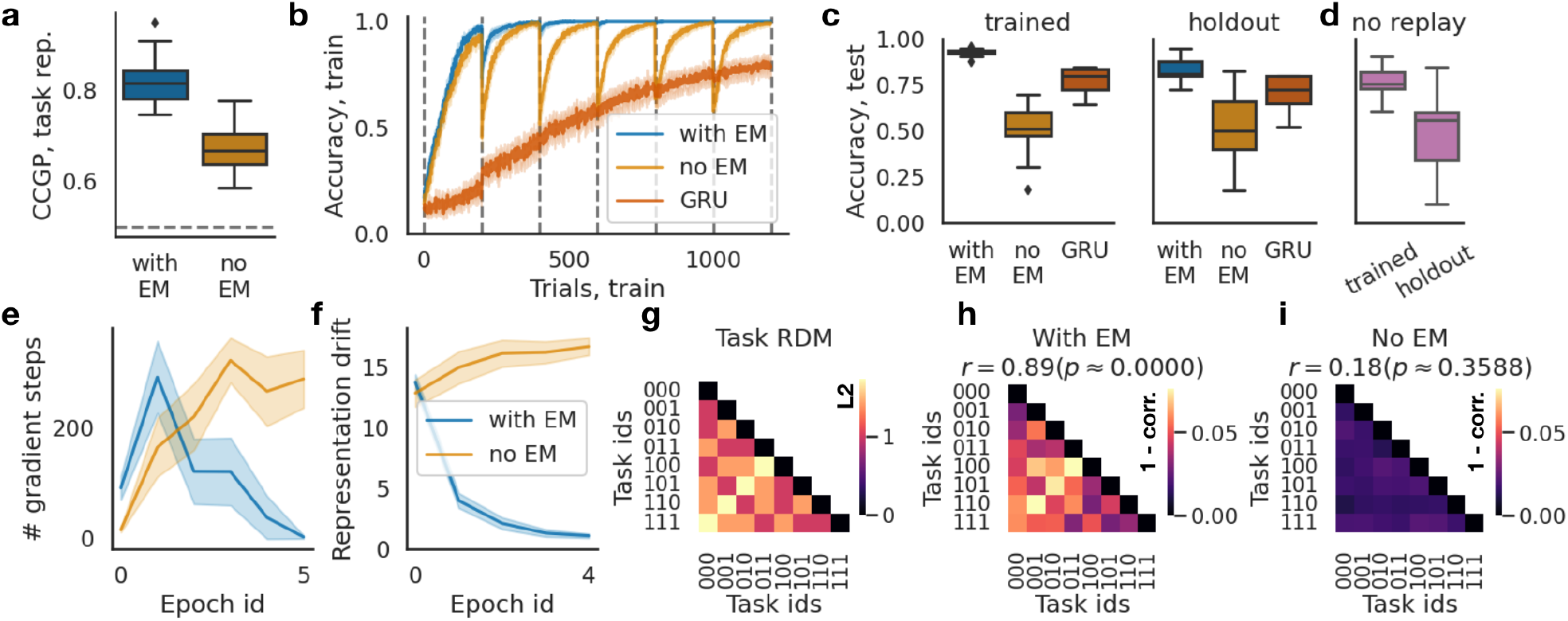
**(a)** CCGP for the task representations is higher for models with EM. **(b)** Prediction accuracy averaged across all seven trained tasks over six training epochs. The dashed lines marked the epoch boundary. GRUs suffered from interference across tasks, slowing learning. No-EM models showed good learning within task blocks but large decreases in performance when the task switched. By contrast, the performance of the with-EM models reached the ceiling more rapidly, as they can retrieve the previously formed task representation. **(c)** During test, accuracy was measured for all eight tasks. The with-EM model was the best at both the trained tasks and the novel holdout task. **(d)** Turning off end-of-sequence replay for the with-EM model reduces performance. This suggests that replaying the just-experienced sequence allowed the with-EM model to encode a more informative task representation. **(e, f)** Models with EM needed fewer gradient steps to find a locally optimal task representation **(e)**, and they had lower representation drift **(f)**, defined as the change of task representation when encountering the same task twice across two consecutive epochs. **(g)** The task RDM computed as the L2 distance between TISs. **(h, i)** The RDMs for task representations during test. The with-EM model RDM **(h)** was highly correlated with the task RDM (Spearman *r* = 0.89, *p <* 10^−4^), but the no-EM model RDM was not **(i)**. N = 30 models. Errorbands indicate 2 SEs.

During the test phase, while the weights were frozen, the model with EM performed the best both for the trained tasks and the holdout task (Fig. 3c). The model with EM also had better CCGP compared to the no-EM model and the GRU (Fig. 3a). These findings of good holdout performance and high CCGP both support the claim that EM promotes structure learning. Additionally, we found that end-of-event replay was important for task performance – when replay was turned off from the with-EM model (Fig. 3d), the performance for both trained tasks and the holdout task was reduced.

We also found that models with EM were more efficient at building the ongoing task representations: They i) needed fewer gradient steps for task inference (Fig. 3e) and ii) had lower representational drift (operationalized as the change of representation when encountering the same task twice) compared to the no-EM model (Fig. 3f).

Finally, to further understand the learned task representations, we compared the representational dissimilarity matrices (RDMs; Kriegeskorte, Mur, & Bandettini, 2008) between the ground truth (Fig. 3g) versus the model representations (Fig. 3h, 3i). Consistent with the CCGP results, the with-EM model RDM was highly correlated with the task RDM (Fig. 3h), and this was not the case for the no-EM model (Fig. 3i). Moreover, this RDM analysis shows that, for the no-EM model, the representations for all tasks became highly similar, which suggests that – by the end of training – the no-EM model had largely “given up” on using the task layer, instead relying on the RNN to support task-specific responding.

To summarize: Without EM, task representations for the same task can drift perpetually. This suggests that finding a task representation according to the ongoing task objective is underdetermined – there are multiple task representations that satisfy the objective. As a result, the task inference process cannot consistently re-discover previously used representations. This is problematic, as having stable task representations is essential for accumulating knowledge about the temporal structure of the ongoing task as well as the relation across tasks. We found that having EM helped to reduce representation drift; effectively, EM acted as a constraint for the underdetermined task inference problem by requiring the model to re-use previously formed task representations.

## Simulation 2. Learning similarity structure

The model’s goal in Simulation 2 was to learn a set of classification tasks, where the tasks differed only in the angle of their decision boundaries (Fig. 4a). The angles ranged from 0 to 360 degrees at 30-degree intervals. Therefore, the relation across tasks was a ring structure (Fig. 4a). At time t, the model received a coordinate *x*, and it had to predict the associated label *y*. As there was no temporal structure in this simulation, we used a feedforward version of our model (the model in Fig. 1a without recurrent connections).

**Figure 4:**
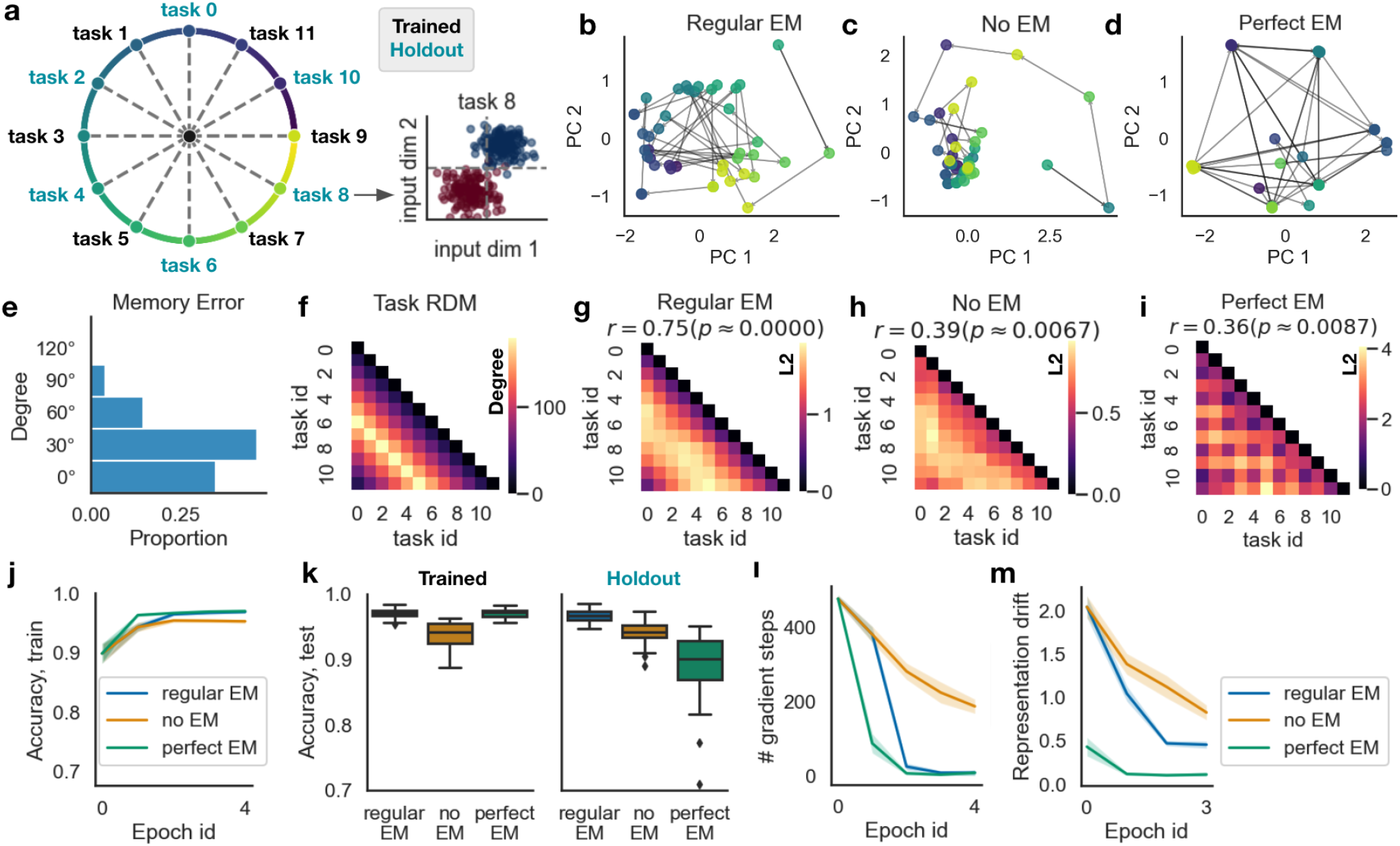
**(a)** The model was given twelve different classification tasks (6 trained, 6 held out), where the ground truth structure across tasks forms a ring. For example, the 8th task is a classification task where the true decision boundary is 240 degrees (counterclockwise from task 0 position). In every epoch, the models experienced tasks with odd indexes in a blocked curriculum (i.e., the trained tasks were 1, 3, 5, …, 11). After training, all models were tested on all 12 tasks, including the holdout tasks (i.e., 0, 2, 4, …, 10). **(b, c, d)** The task representations over blocks during training for the model with EM **(b)** formed a ring structure resembling the ground truth. This was not the case for the model without EM **(c)**. This EM-related benefit was largely due to memory errors, as models with perfectly accurate EM **(d)** also did not acquire the ground truth structure. **(e)** The distribution of memory errors during the test phase shows that similar tasks are more confusable. **(f)** The ground truth RDM that encodes the ring structure. **(g)** The average (over models) RDM for models with regular EM closely resembles the ground truth (*r* = 0.75, *p <* 10^−4^). **(h, i)** The RDM for no-EM models **(h)** and perfect EM models **(i)** had lower correlations with the ground truth. **(j)** The mean performance across all trained tasks during training. **(k)** The mean performance for all tasks during test. Models with regular EM were much better when generalizing to novel holdout tasks. **(l)** Models with no EM needed more gradient steps to find a locally optimal task representation during training. **(m)** Models with no EM showed much higher representational drift for the same task. N = 30 models. Errorbands indicate 2 SEs.

All models were trained for five epochs. In every epoch, the model experienced the odd-numbered tasks (1, 3, 5, …, 11) in a blocked manner. Every block contained 300 trials. The order of the six tasks was randomized. Then, during the test phase, all models were tested on all 12 tasks. The weights were frozen, but the task representation (the purple task layer in 1a) was allowed to be adjusted normally. Note that tasks 0, 2, 4, …, and 10 were holdout tasks, so they were novel to the model during the test phase.

For the regular EM models, memories were indexed by observations, or *{x, y}* pairs. At the beginning of every new task, the model collected a small sample of 10 observations, 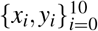. For every observation *{x*_*i*_, *y*_*i*_*}*, the model retrieved the task representation with the closest observation, then returned the average of these task representations. This retrieved pattern was used to initialize the task inference process, in which the task representation was optimized to minimize the loss over 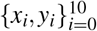 until convergence. These 10 observations were also used as memory keys to be encoded along with the current task representation. In addition to the regular EM model, we also included a *perfect EM* model that used the true task ID as the memory key and consequently never made memory errors (unlike the regular EM model, which sometimes retrieved memories from the wrong task).

We visualized the task representations discovered for every block by projecting them onto the subspace spanned by the first two principal components (PC; Fig. 4b, 4c, 4d). For the model with EM, the task representations formed a ring structure that resembles the ground truth (Fig. 4b). This was not the case for the model without EM (Fig. 4c). Interestingly, this was also not the case for the perfect EM model (Fig. 4d). The finding that regular EM but not perfect EM learned the ring structure shows that naturally-occurring memory errors can help to reveal the similarity structure across tasks. For example, if the ongoing task is task 8, the model might retrieve EM of task representation formed for task 7 or 9 (Fig. 4e), as the observations of these tasks are similar. As a result, representations for these three tasks will be used in overlapping sets of scenarios, making them more similar.

To quantitatively compare the structure across learned task representations versus the ground truth during the test phase, we computed the RDMs (Kriegeskorte et al., 2008) for the ground truth relation across tasks (Fig. 4f), models with regular EM (Fig. 4g), models with no EM (Fig. 4h), and models with perfect EM (Fig. 4i). The model RDMs were averaged across models. We then measured the Spearman correlation between each model RDM versus the ground truth task RDM – the results show that the regular EM model was much better at learning the ground truth structure than the no-EM model and the perfect EM model.

Consistent with the previous simulation, learning the ring structure was useful for generalizing to novel tasks consistent with the learned structure. We tracked the performance for all tasks during training (Fig. 4j) and test (Fig. 4k). Importantly, during the test phase, models with regular EM were much better at generalizing to unseen holdout tasks compared to the two alternative models (Fig. 4k).

Also consistent with the previous simulation, models with EM needed fewer gradient steps for task inference (Fig. 4l). Furthermore, representational drift – the change of task representation when revisiting the same task (Fig. 4m) – was much smaller for models with EM. Without EM, representational drift was high (Fig. 4m) even after task performance converged during training (Fig. 4j). This comparison again suggests that task inference is underdetermined, in the sense that there exist many locally optimal task representations for the ongoing task objective; as such, previously discovered task representations cannot be consistently re-discovered. The role of EM is to constrain task inference by biasing the model to re-use previously found task representations. Importantly, though, the results from the perfect EM condition show that stability alone was not sufficient to drive structure learning, as perfect EM minimized drift but did not permit learning of the ring structure – in this case, the model needed the extra factor of memory errors (present in the regular EM condition) to help stitch the task representations together.

## Discussion

We present a neural network model of how EM can support online inference of task representation and shape the relational structure of task representations. Model comparison results suggest EM played a key role in acquiring the compositional structure in Simulation 1 and the similarity structure in Simulation 2; this, in turn, supported generalization to novel tasks that are consistent with the learned task structure.

Our simulation results indicate two ways in which EM can facilitate the acquisition of structured task representations. First, EM reduces representation drift and stabilizes task representations, which is essential for accumulating knowledge about the relations between tasks. Across the two simulations, we found that, without EM, representations for the same task can drift substantially even after performance converges. This indicates the ongoing task objective is often not constrained enough in the sense that there are many locally optimal task representations. Importantly, not all of these representations reflect how the ongoing task relates to other tasks. Our results show that EM can act as a constraint for task inference by biasing the model to re-use previously discovered task representations.

Simulation 2 also indicates a second, distinct way in which EM can benefit structure learning: When the similarity structure of memory keys is consistent with the similarity structure across tasks, memory errors can help, rather than hinder, the acquisition of the structure. In this situation, memory errors (retrieving a similar task in place of the correct one) will bias the model to use similar representations for similar tasks.

It is notable that perfect (i.e., error-free) EM was sufficient for structure learning in Simulation 1 but not in Simulation 2. We think this is a consequence of the Simulation 1 task being more difficult. That is, in Simulation 1, the model was strongly incentivized to re-use parts of previously learned task representations when learning new tasks because the state transition structure is hard to learn (so “jump-starting” the process with a useful task representation boosts performance a lot). In Simulation 2, the task itself is simple so re-use did not provide as strong of a benefit; in this case, having the extra factor of similarity-based memory errors was helpful for learning similarity structure.

With regard to replay: Our results are consistent with findings showing that humans rapidly replay the just-experienced sequence right before event boundaries (Sols, DuBrow, Davachi, & Fuentemilla, 2017; Silva, Baldassano, & Fuentemilla, 2019) – in Simulation 1, end-of-event replays allowed our model to infer (and then store in EM) a more informative task representation, compared to the task representation inferred based only on data from the start of the task. This predicts that disruption of end-of-event replay would reduce the informativeness of the encoded task representation for later parts of the event but not for the beginning.

In the future, it will be useful to compare our model to other accounts of the role of EM in task representation. Giallanza, Campbell, and Cohen (2023) proposed the episodic generalization and optimization (EGO) framework, which also uses EM to store and retrieve task representations. A key difference is that our model uses EM to initialize task inference, whereas EGO uses backpropagation (through a differentiable EM) to adjust how past memories influence the ongoing task representation. Future work can explore the similarities and differences in the predictions made by EGO versus our model and test them empirically. If the use of EM (shared by the two models) is the key to acquiring structured task representations, then one might expect structured task representations to emerge in EGO in the scenarios described here.

## Acknowledgement

We would like to thank Tyler Giallanza, Declan Campbell, Cody Dong, Jonathan Cohen, and anonymous reviewers for their insightful and constructive feedback.

This work is supported by a Multi-University Research Initiative grant to K.A.N. (ONR/DoD N00014-17-1-2961), an Alan Kanzer Postdoctoral Fellowship to Q.L., and Gatsby Foundation (GAT3708).

